# Integrating animal tracking data into spatial conservation prioritisation for seabirds during their breeding season

**DOI:** 10.1101/2023.12.14.571606

**Authors:** Ruben Venegas-Li, Andre Chiaradia, Harley Schinagl, Akiko Kato, Yan Ropert-Coudert, Hugh Possingham, Richard D. Reina

## Abstract

Understanding the spatial-temporal marine habits is crucial to conserving air-breathing marine animals that breed on islands and forage at sea. This study, focusing on little penguins from Phillip Island, Australia, employed tracking data to identify vital foraging areas during breeding season. Long-term data from sub-colonies and breeding stages were analysed using 50%, 75%, and 90% kernel utilisation distributions (KUDs). Breeding success, classified as low, average, or high, guided the exploration of site, year, and breeding stage-specific habitats. Using Marxan, a widely used conservation planning tool, the study proposes both static and dynamic spatial-temporal scenarios for protection based on KUDs. The dynamic approach, requiring less space than the static strategy, was more efficient and likely more acceptable to stakeholders. The study underscores the need for comprehensive data in conservation plans, as relying on one nesting site’s data might miss essential foraging areas for penguins in other locations. This study demonstrates the efficacy of animal tracking data in spatial conservation prioritisation and marine spatial planning. The dynamic areas frequented emerged as a strategy to safeguard core regions at sea, offering insights to improve the conservation of iconic species like little penguins and promoting the health of islands and the entire marine ecosystem.

## Introduction

Seals, sea turtles, and seabirds, evolved to take advantage of the abundant food supply at sea while still returning to land to breed. Hence, during breeding, these marine animals are exposed to both ocean and land-based threats, including overfishing, bycatch, climate change, pollution, human disturbance, habitat degradation, and alien invasive predators ^1–6^. These threats have increased worldwide ^7,8^, with cumulative impacts on the populations of these marine species ^9,10^. As a result, numerous species within these iconic groups are threatened with extinction ^1,11–13^. Given that these marine animals spend most of their time at sea foraging or resting during their breeding seasons (Hays et al. 2002; Houghton et al. 2002),their conservation should actively protect them in these areas and not focus solely on the land during these crucial periods. However, planning conservation actions for at-sea areas can be complex as there is limited knowledge of how animals use this space^16^.

The challenge of understanding how marine animals navigate their habitats is increasingly being overcome by tracking them through bio-logging, which needs to be leveraged for conservation planning approaches ^17,18^. A clear path forward is incorporating information on habitat use obtained from tracking into planning tools such as spatial conservation prioritisation ^19–21^, a method used to identify areas where management actions can achieve conservation targets efficiently ^22,23^. A common and significant challenge in spatial prioritisation analysis is that species’ use of their habitat is poorly understood. Thus, areas identified for conservation actions are not necessarily directed to those critical zones used by animals, resulting in the inefficient use of conservation resources ^24–26^. Integrating animal tracking data into spatial prioritisation can help to account for differences in habitat use throughout the life cycle of a species according to environmental conditions, to obtain more efficient conservation plans.

Here, using as a case study a mega-colony of little penguins *Eudyptula minor* breeding on Phillip Island, Australia, we explore the inclusion of habitat use information derived from tracking data in the design of management areas for seabirds during their breeding season. We illustrate how different interpretations of the tracking data can affect the efficiency of a conservation prioritisation analysis. Seabirds are central place foragers during their breeding season, i.e., they breed within a place from which they repeatedly depart to forage. Little penguins in Phillip Island breed between August and February. Multi-year tracking data has shown that foraging trips vary in length across three breeding stages, incubation, guard, and post-guard ^27,28^. Furthermore, their use of habitat varies according to environmental conditions. Penguins make short trips, take advantage of the presence of thermoclines, and have a reduced foraging zone in years of high breeding success. In contrast, penguins travel longer distances in an expanded foraging zone in low breeding success years when thermocline is mainly absent ^28–32^. As central place foragers, little penguins that nest in different areas within the colony (sub-colonies) forage in different areas, e.g., there is little overlap between the core foraging areas of sub-colonies that are only 2 km apart ^28,33^.

This study addresses the following research questions for central place foragers: i) how much more efficient could a dynamic conservation plan be compared to a static conservation plan for Phillip Island’s little penguins; ii) do we need data from different years and breeding cycles to plan efficient and effective conservation areas for central place foragers?; iii) do we need data from different breeding sub-colonies (or that are distributed throughout the colony) to plan efficient and effective conservation areas for central place foragers?

## Results

### Little penguins foraging areas

The tracking data of the little penguins shows that: i) individuals nesting in the Penguin Parade site forage in different locations than those nesting in Radio Tracking Bay (Fig 1b), ii) foraging areas during the guard period are smaller than during the other 2 breeding stages and tend to be smaller during years of high breeding success (Fig 1a).

**Figure 1.**
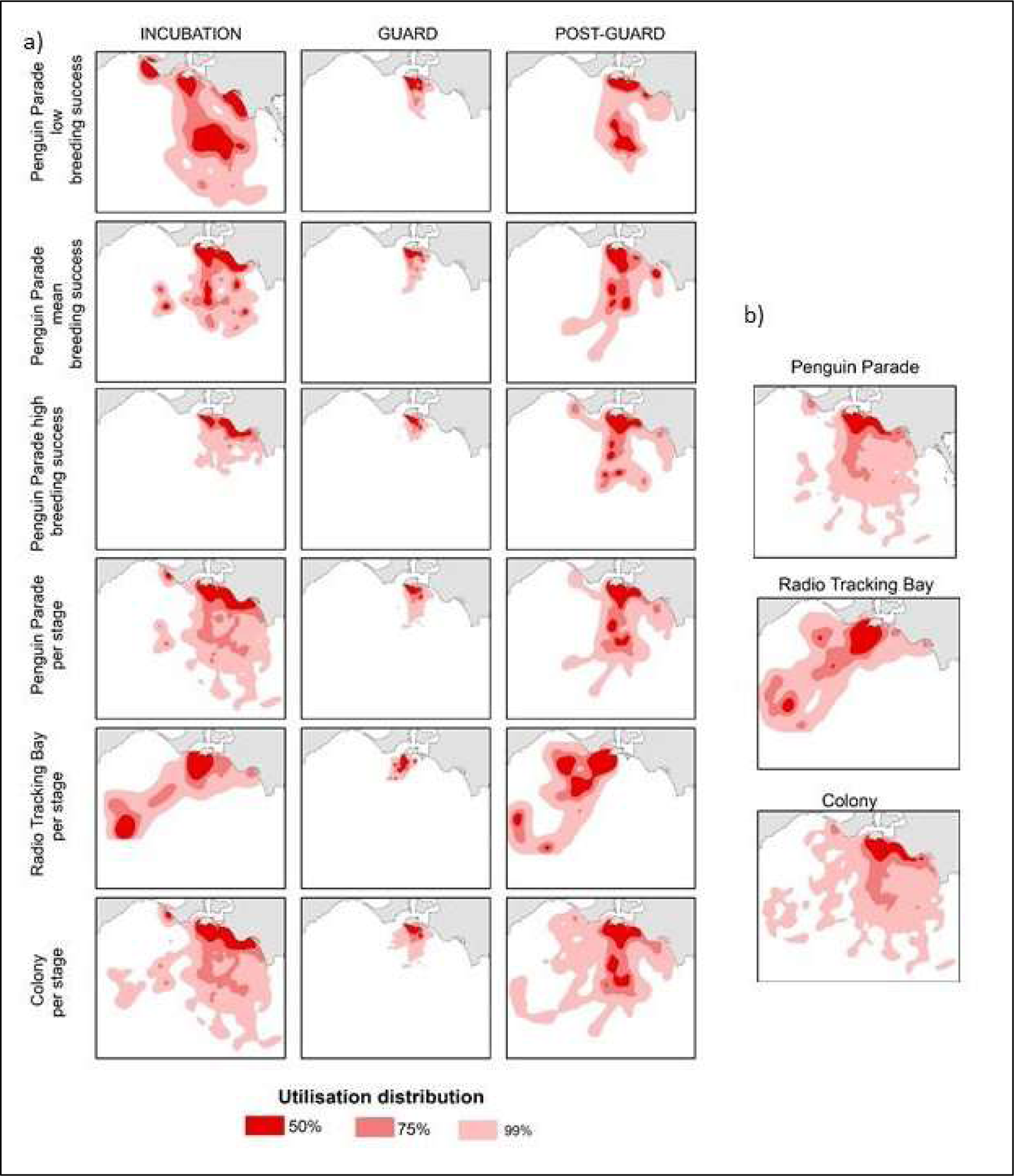
Utilisation distributions of foraging areas used by little penguins from Phillip Island, considering temporal differences within a breeding season (a), and disregarding temporal differences (b).

### Static and Dynamic conservation planning

The static and dynamic prioritisation scenarios representation targets were achieved for all the conservation features, and conservation priorities are clearly defined as shown through the selection frequency maps (Fig 2. scenarios 1 and 2a-2i). On the other hand, through scenario 3, which targeted only the protection of 18.5% of the overall foraging range of the little penguins, as expected, we did not identify conservation priorities. Areas selected for protection as part of the best solution from scenario 3, would (75-99% UD), meaning that the core foraging range of the penguins is underrepresented according to our targets.

**Figure 2.**
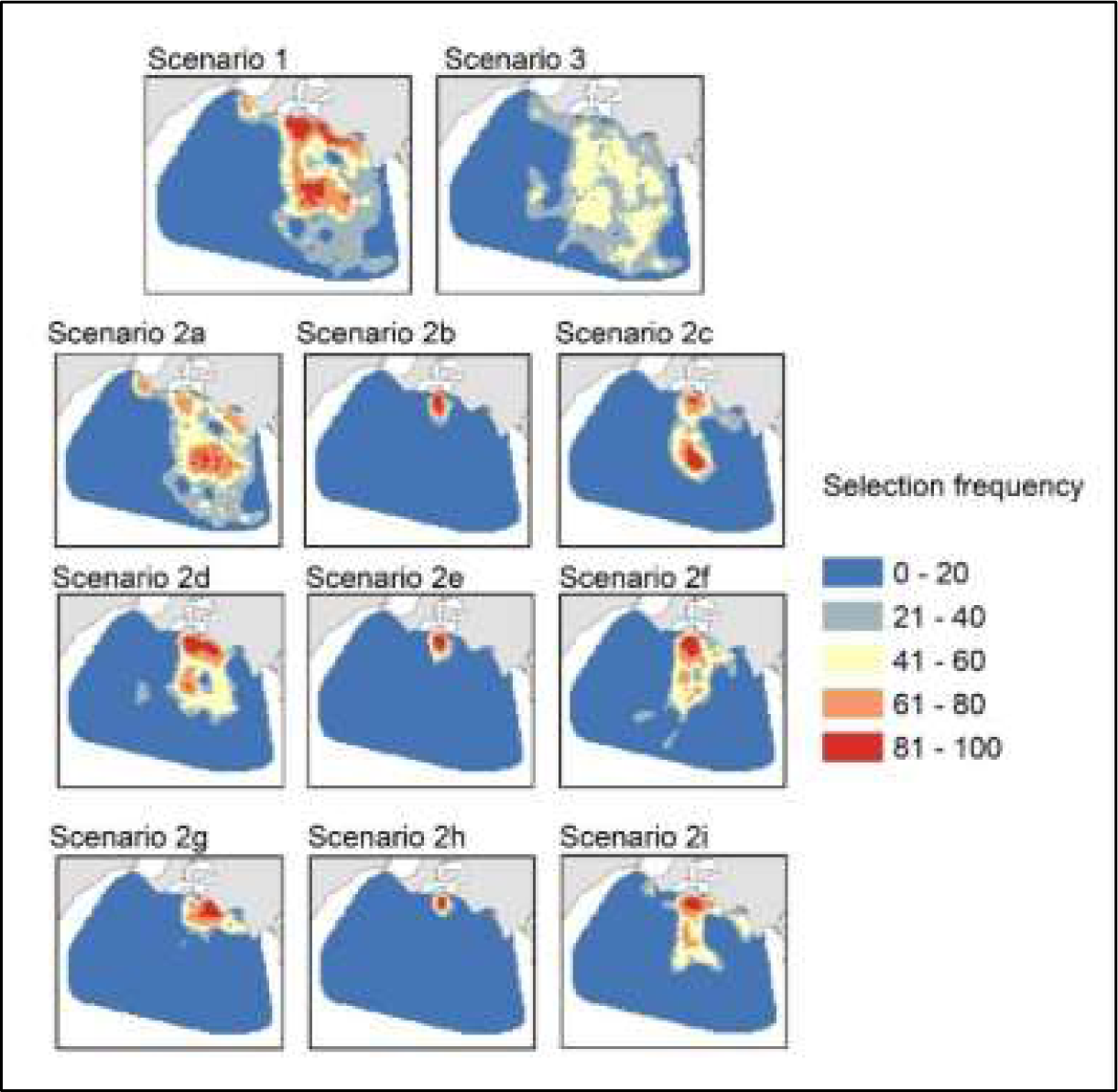
Selection frequencies for scenario 1, 2a-2i and 3.

In the static scenario 1, 18.5% of the planning area was selected (Fig 3). A dynamic conservation plan would have conservation areas that cover approximately half, or less than half, of what is needed in a static conservation scenario at different points in time. The exception is during the incubation period of years with low breeding success, in which the area that needs to be protected is the same as that for the static conservation plan (Figs 2 and 3).

**Figure 3.**
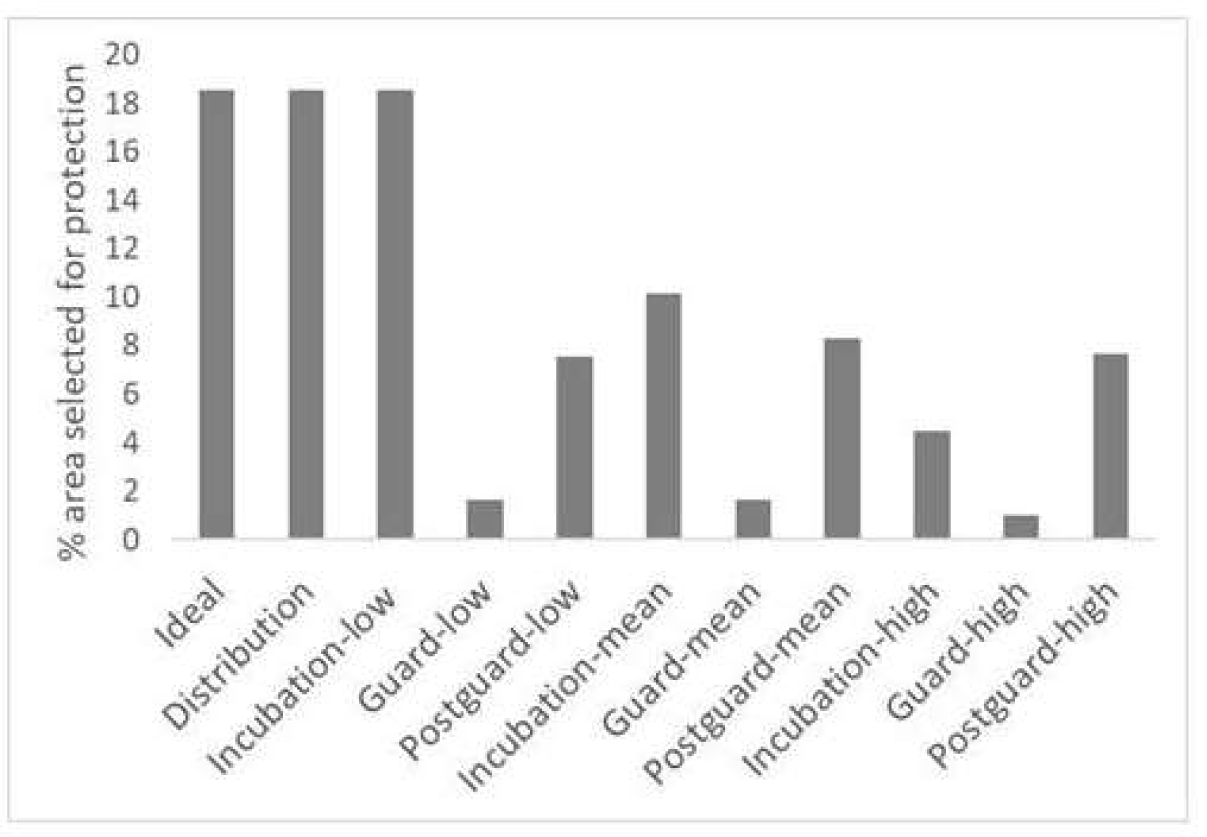
Total planning area (%) that was selected as part of the best solution in scenarios 1, 2a-2i, and 3.

### Conservation planning with a limited understanding of temporal differences in habitat use

Failing to account for temporal differences in habitat use by little penguins in spatial prioritisation analysis resulted in conservation priorities shifting spatially (Fig 4a). More importantly, we show that failing to account for temporal differences results in a selection of sites where conservation targets are not achieved for multiple UDs (conservation features), i.e., solutions are not efficient (Fig 4b). For example, the areas selected for protection in the best solution of scenario 4 will fail to achieve the targets set for 15% of the UDs estimated per stage and per breeding season category. Scenario 4 ignores differences in habitat use between years with different breeding year categories. A complete disregard for temporal differences in habitat use (scenario 5) results in almost 60% of the targets set for the UDs estimated per stage and per breeding year category to be unmet. Mostly, the targets for the 50% and 99% UD are unmet.

**Figure 4.**
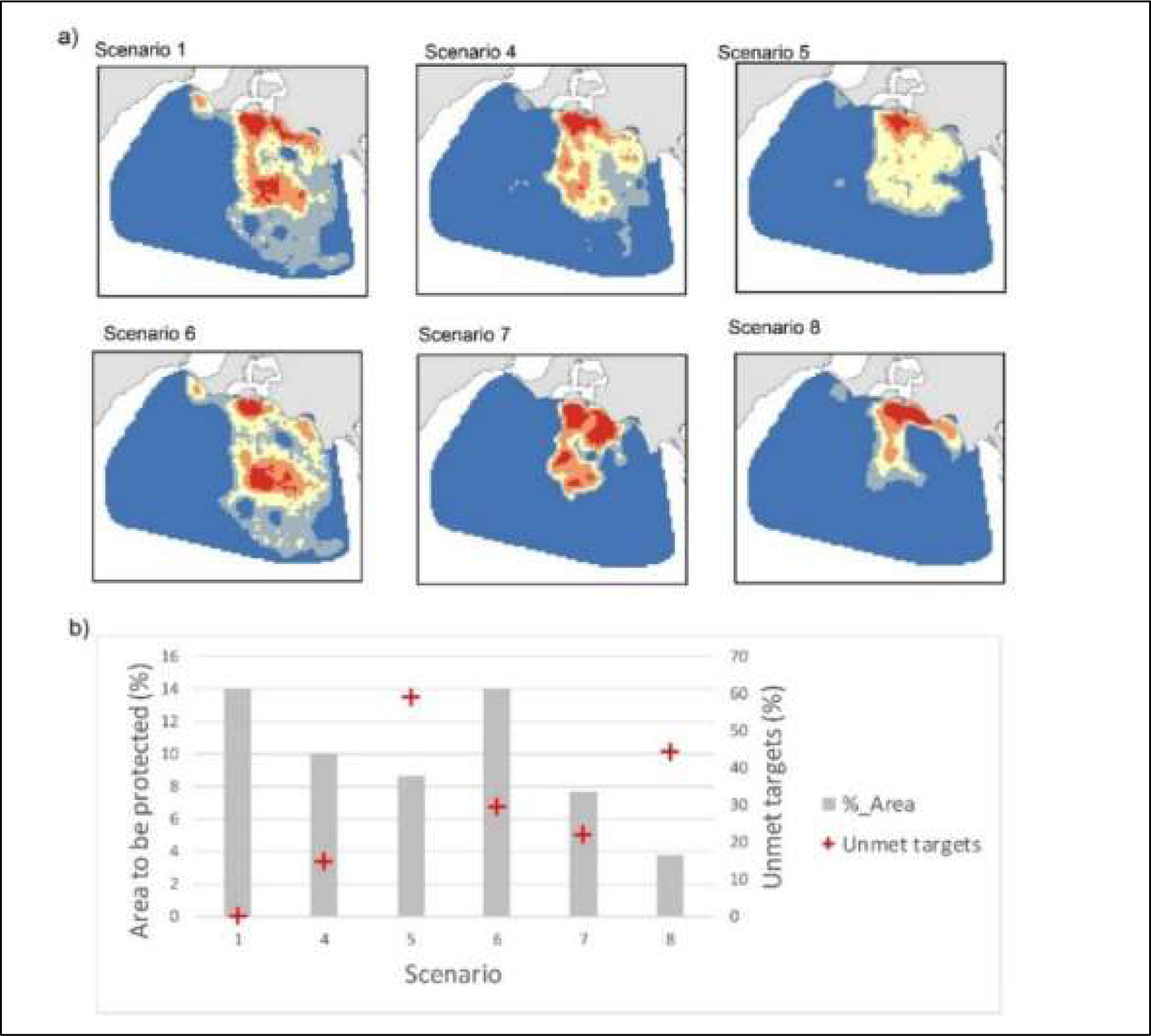
Selection frequencies for scenarios that capture different levels of detail in temporal dynamics of foraging ranges by Phillip Island’s little penguin (a). Total planning area (%) that selected as part of the best solution in each of the scenarios, and number of foraging areas for which representation targets are not reached.

Conservation planning based on data for only one year (equal to one category of breeding success) such as scenarios 6,7, and 8 will also result in unmet conservation features targets (Fig. 4b). For example, using data only for a year with a high breeding success, when penguins forage closer to the coast, will result in over 45% of foraging ranges to be underrepresented in conserved areas.

### Conservation planning with a limited understanding of within-colony differences in habitat use

We show that not accounting for differences in habitat use between the two sub-colonies could result in an inadequate representation of core forging ranges in spatial plans for central-placed foragers. Conservation plans that use tracking data collected only in one sub-colony (Fig 5a, scenarios 10 and 11), would bias protection to those areas where the individuals from each sub-colony forage. It results in the unmet representation targets for the foraging ranges of the sub-colony for which data were not used in the planning (Fig 5b). Likewise, conservation plans will be inefficient if they are generated using data in which UDs are calculated by aggregating all the data available for the Phillip Island penguin colony (scenario 12), without recognising that individuals nesting in different areas in the colony have different foraging areas (Fig 5a, scenario 12). Figure 5 shows that the resulting priorities for scenario 12 are biased towards the foraging areas of penguins nesting in the Penguin Parade, likely because there is more data for that site, with none of the conservation features for Radio Tracking Bay achieving its targets.

**Figure5.**
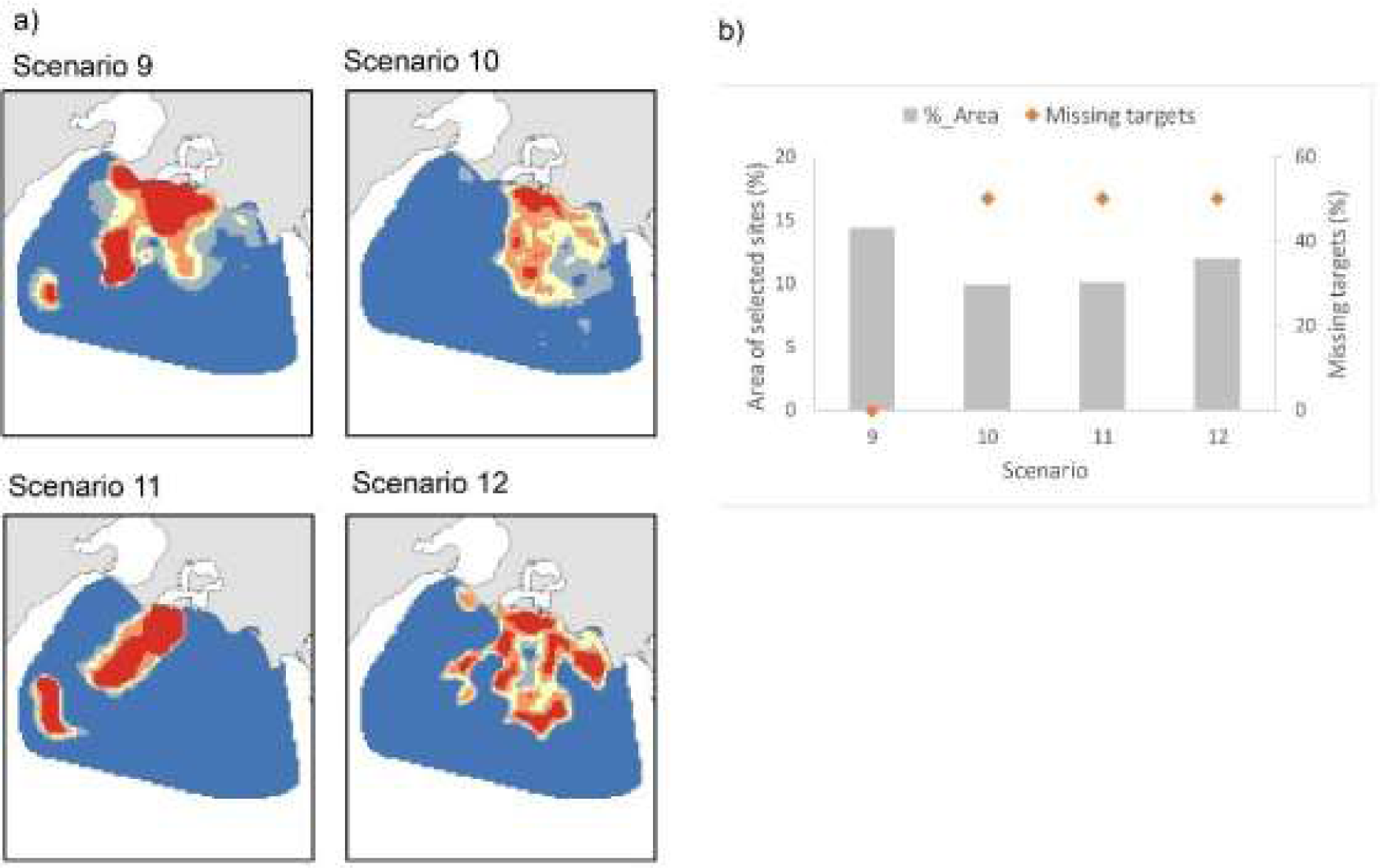
a) Selection frequencies for scenarios that capture different levels of detail in within-colony variability of foraging areas by Phillip Island’s little penguin. b) Total planning area (%) that is selected as part of the best solution in each of the scenarios, and number of foraging areas from an ideal scenario (where within-colony differences in habitat use are captured and use in planning) for which representation targets are not reached.

## Discussion

We investigated an innovative approach to incorporate habitat use information derived from tracking data of a seabird breeding on an island into spatial conservation prioritisation to plan at-sea conservation actions during their breeding seasons. Our approach can be more efficient in targeting conservation actions to areas most used and can be applied to design both static and dynamic spatial plans. However, our results highlight the importance of understanding that limited tracking samples in space and time, and different interpretations of the available data, can produce inefficient or suboptimal solutions. Thus, we emphasize the value that animal tracking data has for understanding how animals use their habitat, and hence how valuable this can be for spatial conservation planning. In the marine environment, particularly on islands, where the direct and continuous observation of individuals is difficult, we showed that tracking data can be fundamental to understanding their behaviour and planning their conservation. Our study contributes to the much-needed ^17,41–44^ and growing literature exploring how to incorporate tracking data into spatial prioritisation^19–21,45–47^. Our approach could be applied to central place-based foragers with different breeding stages, aggregating during the breeding seasons and with relatively short feeding ranges^48^.

Using tracking data to estimate utilisation distributions for each breeding stage and breeding success class as a proxy for environmental conditions, allowed us to compare static and dynamic spatial conservation plans. Most configurations of the dynamic plans would occupy less than 50% of the area that would be needed through a static plan to achieve the conservation targets. Other studies have found that dynamic management could prove more efficient than static ones ^20,49–51^. In areas where economic activities such as tourism and fishing, dynamic management can better balance ecological, economic, and social objectives (Maxwell et al., 2015), potentially increasing ocean users’ acceptability of marine spatial plans ^52^. Although dynamic ocean management has already been implemented and new tools are under development ^53^, a persistent challenge in our specific case study involves accurately forecasting environmental conditions and penguin breeding success before the onset of the breeding season. The precautionary approach can provide a framework for decision-making based on worst-case scenarios to guide conservation efforts. In marine spatial planning, decision-makers can apply our conservative dynamic area in conjunction with scientific research and stakeholder engagement to make informed decisions. In the case of predicting the area usage at critical stages of little penguin breeding, a precautionary approach could be applied using larger dynamic areas based on years of low breeding success.

A key assumption in this study is for all foraging areas, which change across breeding stages and years, to be protected for conservation plans to be effective. While tracking data can prove key for doing this, our results show how different interpretations of the data could affect the results of spatial prioritisation analysis. Scenarios that aggregate all the data to estimate UDs ignoring temporal differences in habitat use fail to protect many of the foraging areas. When differences in habitat use between individuals nesting at different sub-colonies are ignored, UDs were estimated on a per colony basis instead of per sub-colony basis.

Moreover, our results also demonstrate that limited data availability can hinder the efficiency and effectiveness of spatial plans created using tracking data. For example, scenarios that use data limited temporally, e.g., data which only captures spatial usage during one season, will fail to reach the conservation targets for the core foraging areas in seasons with different environmental conditions. Additionally, we show that collecting tracking data from only one sub-colony and planning based on these data, will result in conservation plans that will fail to protect the core foraging areas of penguins from other sub-colonies. This result supports the suggestion by Sanchez et al. ^28^ for tracking studies to best capture the foraging behaviour within a colony to inform conservation plans. They suggest that for large seabird colonies, tracking data should be collected through different breeding stages and years, and from more than one breeding site ^54^.

The optimal design of marine spatial plans depends on information and knowledge of how anthropogenic impacts interact with the species that are mobile in the system. However, challenges in surveying mobile species over adequate spatial and temporal scales can mask the importance of particular habitats, leading to uncertainty about areas to protect to optimize conservation efforts. Thus, the incorporation of spatial and temporal animal tracking data into marine spatial planning is crucial for understanding the habitat use and behaviour of marine animals. The collection of tracking data, especially for central-place foragers with different breeding stages, across various breeding sites, and over multiple years creates dynamic areas that increase certainty about areas to protect, as shown in this study. Our data on dynamic areas can assist in prioritizing conservation actions based on habitat use data derived from tracking of flagship species in the marine system to ensure preservation of critical foraging hotspots and increasing the effectiveness of spatial plans. Flagship species like little penguins are vital for conserving the entire marine ecosystem, reflecting the well-being of numerous other species and habitats in our oceans. This foundational knowledge can guide the implementation of conservation strategies, safeguarding the long-term health of marine ecosystems. Our findings highlight the importance of leveraging animal tracking data into management strategies that can enhance the effectiveness of marine spatial planning.

Animal tracking data provides the opportunity to acquire a wealth of information about where and when marine animals move and how they interact with their environment. This study emphasises the pivotal role of such data in the context of islands, where marine animals, like little penguins, exhibit intricate movement patterns tied to their breeding cycles and marine environmental conditions. The significance of this understanding becomes evident in formulating effective conservation strategies that specifically target crucial foraging areas, recognising the variability within and between breeding seasons. Failing to do so, could result in conservation actions that perform poorly, as occurs with Marine Protected Areas in protecting shy Albatross in the Bass Strait^26^. Our results showed that for little penguins, examining animal movement through the different breeding stages and years would help to create dynamic spatial plans that are more efficient than static plans. It also highlighted that limited tracking samples in space and time, and different interpretations of the available data can affect the results of spatial prioritisation analysis to produce inefficient or suboptimal solutions. Our study contributes to the broader understanding of how animal tracking data can inform conservation planning in island ecosystems, emphasising the need for a nuanced and dynamic approach to safeguarding marine biodiversity.

## Methods

Phillip Island (38.5065° S, 145.1496° E, Fig 1) is home to a little penguin population of over 32,000 individuals, the largest in the world, that has doubled since the 1990s, due to conservation actions taken on land ^34^.

### Inferring habitat use from penguin tracking data

We tracked 237 little penguins from two sub-colonies using GPS loggers. The two sub-colonies, named Penguin Parade and Radio-Tracking Bay are separated by 2 km (Fig 1). In total, 373 foraging trips over incubation (n = 84), guard (n = 193; chicks between 1 and 19 days old), and post-guard (n = 96; chicks are unattended during the daytime in the colony) were analysed. At the breeding stage, birds from both sub-colonies were tracked simultaneously, aiming to maintain the same sample size per site and sex. Birds were captured in their nest and immediately returned after the tracking device attachment, with a handling time of less than five minutes. Tracked individuals were recaptured in their nests after a single foraging trip to retrieve the loggers. The tracking device deployment procedure was moderate with the stress of short duration and brief restraint for fitting devices to wild animals. This procedure was performed in accordance with relevant guidelines and regulations under the Australian Code for the Responsible Conduct of Research, with approval from the Phillip Island Nature Parks Animal Ethics Committee and a research permit issued by the Victorian Department of Environment, Land, Water, and Planning, Australia.

We used the little penguin tracking data to estimate how much they used different marine areas for foraging by calculating the 50%, 75%, and 99% utilisation distributions (UD) through kernel density analysis. To calculate these, we used the kernelUD() function from the “adehabitatHR” R package ^35^. The smoothing parameter h was calculated using the ad hoc method ^36^. The utilisation distributions were calculated using different groupings of the tracking data (Table 1). For example, we calculated UDs for each sub-colony separately, and for the entire colony (using data for both sub-colonies, i.e., disregarding differences in the use of foraging areas by penguins nesting in the two sub-colonies). We also aggregated the data for each site, either ignoring intra-annual or interannual differences in habitat use. The different groupings allowed us to build a static and dynamic spatial conservation prioritisation approach, and to test how data availability or different interpretations of the data would affect conservation plans.

**Table 1.**
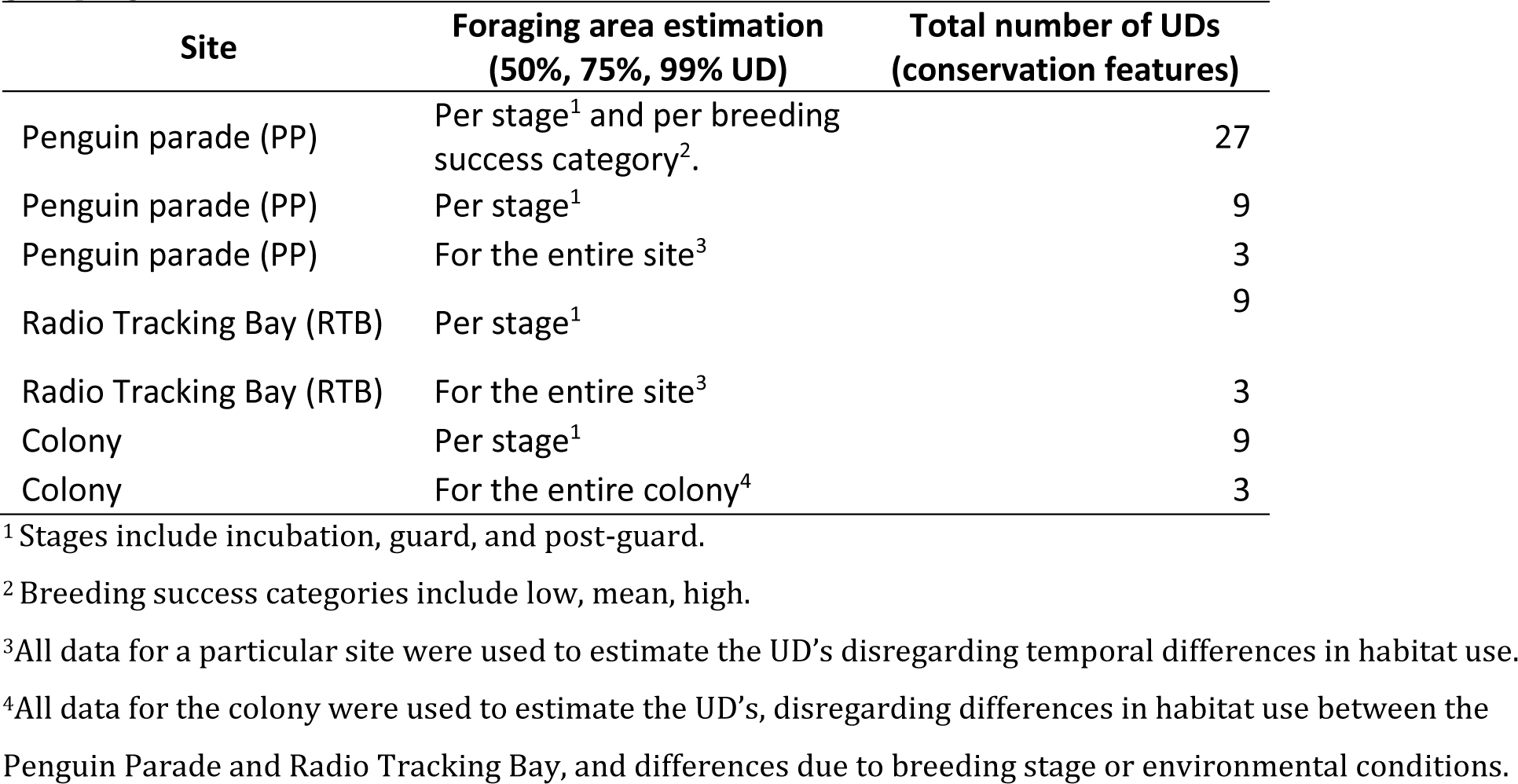
Kernel utilisation distributions estimated from tracking data as a measure of habitat use by little penguins from two different nesting sub-colonies in Phillip Island, Australia, using different <u>groupings of data.</u>.

### Spatial prioritisation

Little penguins’ foraging area varies across breeding stages and over years ^28,30,31,33^. This variability suggests that efficient and effective conservation plans for the little penguins should target the different areas where individuals from the colony forage, during all breeding stages and under different environmental conditions. Conservation plans based on penguin’s tracking data from years with high breeding success when they stay closer to the coast could overlook and not protect crucial foraging areas in years of low breeding success, when they would forage further away from the coast.

Thus, our planning objective was to select sites within our planning region (penguin foraging area) where representation targets for the penguin’s feeding areas are achieved, based on how much and when they are used (utilisation distributions). We set arbitrary conservation targets of 70%, 50%, and 30% for the 50%, 75% and 99% UD, respectively. From this point forward, the utilisation distributions will be referred to interchangeably as conservation features.

We defined the overall penguin’s foraging area (planning region) as the smallest convex polygon enclosing all the different UDs. This foraging area was subdivided into a total of 1979 planning units of 4 km^2^, although some of these were smaller as they were intersected by the study region’s boundaries. To identify sets of planning units where our targets are met efficiently, we used the spatial prioritisation software Marxan ^37^. Marxan uses a simulated annealing algorithm ^38^ to identify near-optimal site configurations in a study region where defined quantitative conservation targets can be achieved while minimising cost. We assigned a uniform cost to all planning units, to focus on differences between the solutions based on the tracking data. We calibrated the boundary length modifier (BLM) in Marxan to find compact solutions following Stewart and Possingham ^39^.We ran Marxan 100 times with 1,000,000 iterations per run for each planning scenario, potentially producing 100 different solutions each time. To evaluate the results of the different scenarios, we used the best solution, and the selection frequency outputs from Marxan. The best solution is the run with the lowest Marxan score, arguably the most efficient solution. The selection frequency is the number of times a particular planning unit is selected across all the runs and indicates how important a planning unit is to achieve efficient solutions in a particular scenario.

### Using tracking data in spatial prioritisation to plan static and dynamic conservation plans

We designed a first set of planning scenarios to demonstrate how to incorporate habitat use information obtained from animal tracking data into spatial prioritisation. We only used data for the Penguin Parade site for this part of the study, as data for Radio Tracking Bay was available for only one year with average breeding success.

For an “ideal” static planning scenario (scenario 1, Table 2), we set representation targets for each UD calculated for the Penguin Parade per stage and per breeding success category (27 conservation features in total: 3UDs x 3 breeding stages x 3 breeding success categories). With the term static, we mean that the areas selected to carry out a given conservation action (e.g., a marine protected area) would be fixed both in space and time over the breeding seasons and years. The term ideal refers to all foraging areas being adequately represented in conservation areas. On the other hand, in a dynamic planning scenario, areas where conservation actions are to be applied can change in time across breeding stages and according to the environmental conditions of a given year - breeding success is used as a proxy ^40^. Therefore, scenario 2 comprised a set of 9 “sub-scenarios” in which we separately targeted protection for the UDs of each combination of breeding stage and breeding success category.

**Table 2.**
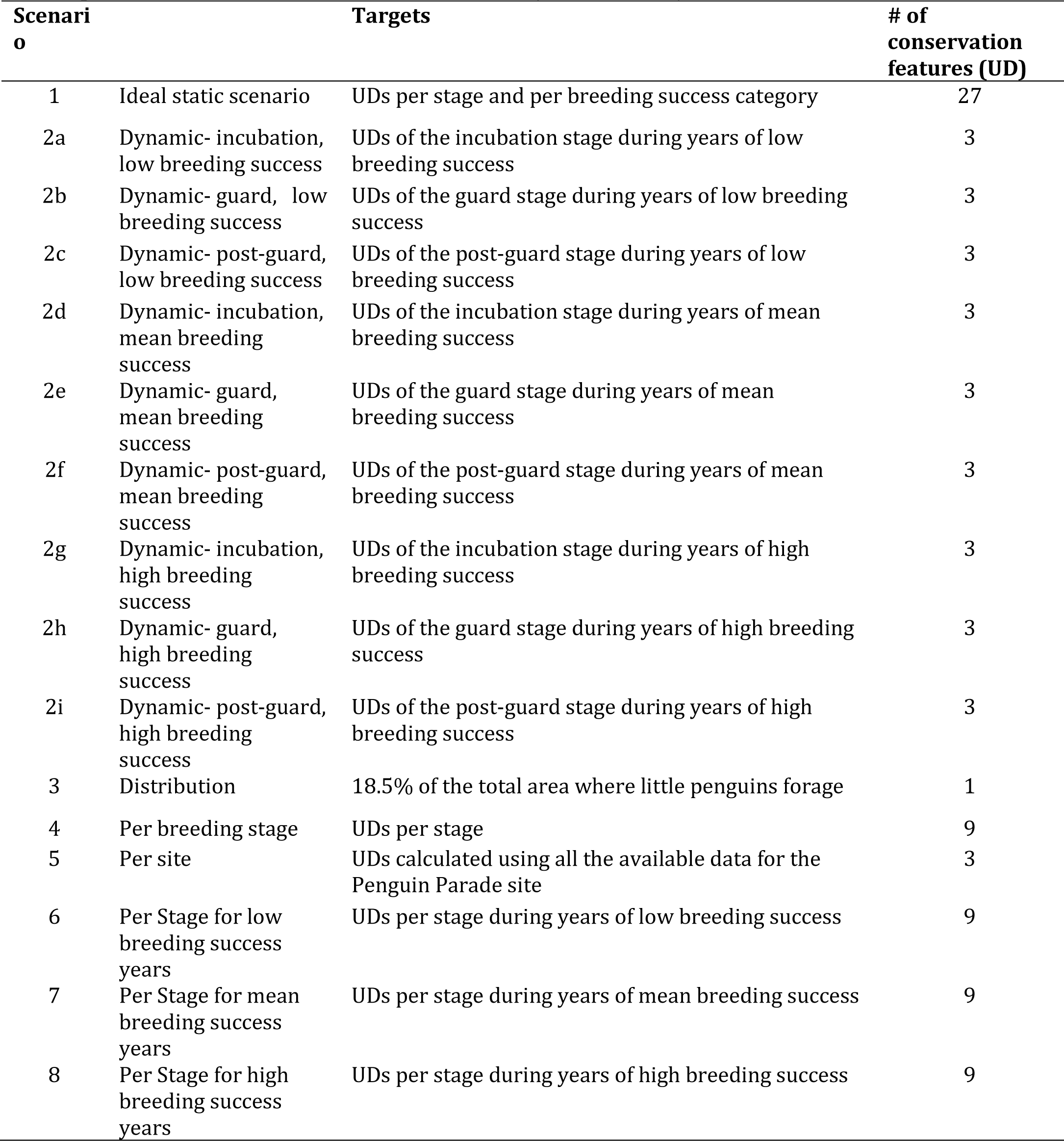
Scenarios using only data from Penguin Parade for demonstrating static and dynamic conservation prioritisation approaches (Scenarios 1-3) and assessing limitations in the temporal dimension of habitat use information (scenarios 4-8).

To illustrate the benefit of scenarios 1 and 2 in protecting the different foraging areas according to how much and when they are used, we designed scenario 3 which targets to protect only a percentage of the broad foraging range of little penguins. This scenario disregards differences in habitat use, as commonly done in spatial prioritisation analysis. We set a representation target of 18.5% for the area where little penguins tracked from the Penguin Parade forage, which equals the total area protected in scenario 1.

### Conservation planning with limited consideration to temporal differences of habitat use

Using data only for the Penguin Parade, we used a second set of planning scenarios to test how conservation plans change if we fail to capture temporal differences in habitat use. Scenarios 1 (Ideal static scenario), 4, and 5 (Table 2) represented scenarios where the same data are interpreted differently, increasingly losing temporal detail. Scenario 4 disregards differences in habitat use between years with different environmental conditions (breeding success categories), and scenario 5 ignores any temporal differences in habitat use.

Furthermore, scenarios 6, 7, and 8 were designed to assess whether conservation plans that used data of only one breeding year (represented by the data for each breeding success category) to inform habitat use, would fail to protect all the feeding areas of the little penguins over the years. This is not an uncommon problem in animal tracking projects, due to the relatively high costs of equipment and tracking resulting in incomplete or patchy data.

To assess how losing temporal detail in habitat use would affect conservation plans, we compared scenarios 4, 5, 6, and 7 to scenario 1. Specifically, we quantified the number of foraging areas estimated for the penguin parade per stage and breeding year’s category, underrepresented (doesn’t achieve conservation targets) in the best solution of those scenarios where temporal detail is lost.

### Conservation planning with limited understanding of within-colony differences in habitat use

We used four prioritisation scenarios (Table 3) to investigate how conservation plans could change if we fail to capture within-colony variability in habitat use. For all scenarios, UDs were estimated per stage for the different aggregations of data for each sub-colony, and not per stage/breeding success category. The reason for this is that data for Radio Tracking Bay was limited to one year with mean breeding success. In scenario 9, we set conservation targets for the UDs estimated for each site per breeding stage separately. Given that this scenario captures both temporal and spatial differences in habitat use, it is the baseline to which the other three scenarios were compared. In scenario 10 we set targets only for the UD calculated for the Penguin Parade site per stage (equal to scenario 4); and in scenario 11 we set targets for the UDs estimated with data from RBT per stage only. Scenarios 10 and 11 assess limitations that could arise from conservation plans created using tracking data information for only one of the sub-colonies. Finally, in scenario 12 we set conservation targets for the UDs calculated by integrating data from both subcolonies per stage, to test how conservation plans could be affected in a case in which within-colony differences in habitat use are ignored, even if the data is available.

**Table 3.**
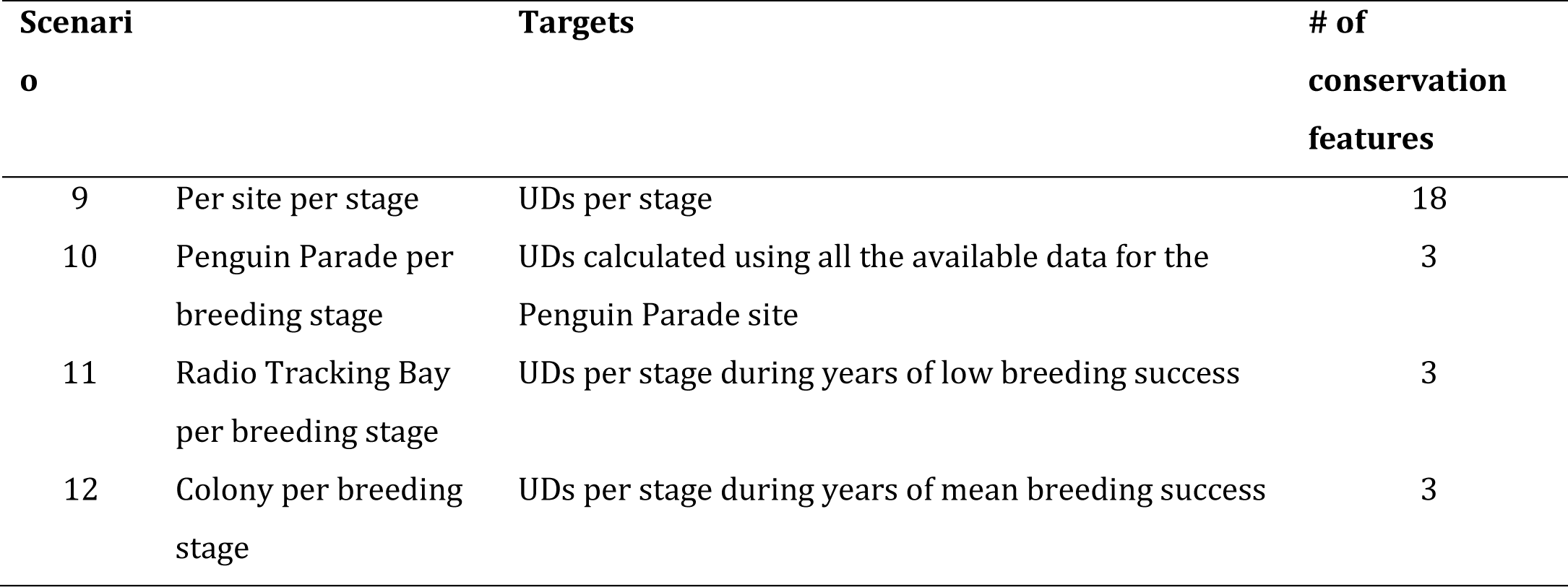
Scenarios used for assessing how spatial conservation plans change when there are limitations in our understanding of within-colony differences in habitat use.

## Acknowledgements

We thank all the students and volunteers for their tireless support in collecting these data over the years, particularly the Nature Parks’ research technical staff Leanne Renwick, Paula Wasiak, Meagan Tucker, Marjolein van Polanen Petel and Jordan Roberts. We thank Phillip Island Nature Parks for continuing support for penguin research.

## Ethics statement

Work was conducted in accordance with Phillip Island Nature Parks Animal Ethics Committee approvals over the years, numbers 2.2004, 3.2007, 2.2010, 2.2013, 3.2013, 3.2016, 4.2019 and 4.2021, and research permits issued by the Victorian Department of Environment, Land, Water, and Planning, Australia, numbers 10003049, 10003419, 10004360, 10005605, 10007320, 10008506, 10010038 and 10010400. All experiments were performed in accordance with relevant guidelines and regulations under the Australian Code for the Responsible Conduct of Research (<u>the Code</u>) which sets out principles and responsibilities that both researchers and institutions to conduct research under the auspices of Australian institutions. The tracking device deployment procedure was moderate with stress of short duration and brief restraint for fitting devices to wild animals.

## Author contributions

RVL, AC, RD and HP conceived the manuscript. AC, AK and YRC performed the fieldwork. AK and YRC processed the tracking data. RVL and HS performed the statistical and GIS analyses. RVL led the writing up with contributions from all authors.

## Additional Information

### Data accessibility

Datasets used for this manuscript are available at: https:// doi. org/ 10. 57745/ IT6ITA.

